# Truncation and Motif Based Pan-Cancer Analysis Highlights Novel Tumor Suppressing Kinases

**DOI:** 10.1101/254813

**Authors:** Andrew M. Hudson, Natalie L. Stephenson, Cynthia Li, Eleanor Trotter, Adam J. Fletcher, Gitta Katona, Patrycja Bieniasz-Krzywiec, Matthew Howell, Chris Wirth, Simon Furney, Crispin J. Miller, John Brognard

**Author notes:** Indicates equal contribution by authors (Andrew M. Hudson, Natalie L. Stephenson).

## Abstract

A major challenge in cancer genomics is identifying driver mutations from the large number of neutral passenger mutations within a given tumor. Here, we utilize motifs critical for kinase activity to functionally filter genomic data to identify driver mutations that would otherwise be lost within mutational noise. In the first step of our screen, we define a putative tumor suppressing kinome by identifying kinases with truncation mutations occurring within or before the kinase domain. We aligned these kinase sequences and, utilizing data from the Cancer Cell Line Encyclopedia and The Cancer Genome Atlas databases, identified amino acids that represent predicted hotspots for loss-of-function mutations. The functional consequences of new LOF mutations were validated and the top 15 hotspot LOF residues were used in a pan-cancer analysis to define the tumor-suppressing kinome. A ranked list revealed MAP2K7 as a candidate tumor suppressor in gastric cancer, despite the mutational frequency of MAP2K7 falling within the mutational noise for this cancer type. The majority of mutations in MAP2K7 abolished catalytic activity compared to the wild type kinase, consistent with a tumor suppressive role for MAP2K7 in gastric cancer. Furthermore, reactivation of the JNK pathway in gastric cancer cells harboring LOF mutations in MAP2K7 or JNK1 suppresses clonogenicity and growth in soft agar, demonstrating the functional importance of inactivating the JNK pathway in gastric cancer. In summary, our data highlights a broadly applicable strategy to identify functional cancer driver mutations leading us to define the JNK pathway as tumor suppressive in gastric cancer.

**Summary:** A unique computational pan-cancer analysis pinpoints novel tumor suppressing kinases, and highlights the power of functional genomics by defining the JNK pathway as tumor suppressive in gastric cancer.

---

It was estimated that by the end of 2017 more than 1.6 million cancer samples would have been sequenced by next generation sequencing (NGS) [1]. The greatest challenge now lies in interpreting this data to dissect tumorigenic mechanisms and identify therapeutic targets. A major problem is that the data is often noisy with many inconsequential passenger mutations obscuring the detection of driver mutations [2–3]. Now that most cancer subtypes have been characterized by large-scale sequencing studies, the common drivers have been identified [4]. However, the fact that many of the samples in these studies do not have an identifiable common driver suggests there are a multitude of lower frequency drivers that we struggle to detect above the noise [5–6]. The best method to discover more cancer drivers is under debate [7–9]. Should we continue sequencing more and more samples, or do we focus on functional studies? Currently, in silico methods are already widely used to attempt functional analysis of large genomic data sets [2,10], however these assessors are limited and may miss functional driver mutations [11–14]. Therefore, there is a need to improve genomic analysis to assist in unlocking the potential of these huge public datasets. By better linking existing knowledge of a proteins function to the associated structural features we can begin to functionally screen genomic data. Protein kinases are a well-characterized class of proteins with documented mechanisms linking structural motifs to protein function [15–17]. This makes them ideal candidates to develop motif-driven bioinformatic screens.

We initially produced a list of candidate tumor suppressing kinases using the frequency of truncating mutations from The Cancer Genome Atlas (TCGA) and the Cancer Cell Line Encyclopedia (CCLE) that would abolish catalytic activity. Sequence alignment of this tumor suppressing kinome allowed the identification of mutational hotspots in conserved regions. The top 12 mutational hotspots were all within motifs already known to be critical for kinase function ‐validating this approach of hotspot LOF mutation identification. Two novel hotspot residues were biochemically analyzed, and found to also result in inactivation of the kinases harboring the mutation. We then developed a bioinformatics screen to identify mutations in the top 15 hotspot residues in 411 canonical kinases in TCGA and CCLE datasets. Kinases were ranked by the frequency of these mutations. Alongside known tumor suppressors such as STK11, we identified and validated a high incidence of MAP2K7 LOF mutations in gastric cancer, and highlight a tumor suppressive role for MAP2K7 and the JNK pathway in this cancer subtype. There has been great debate regarding the role of the JNK pathway in cancer as indicated by the numerous contradictory publications in the literature [18]. The genetic make-up of the tumor and the tumor microenvironment will dictate an oncogenic or tumor suppressive role for this pathway in various cancers [19]. Our study provides a framework to assess the role of kinases in various types of cancers and our results highlight a tumor suppressive role for the JNK pathway in gastric cancer.

## RESULTS

### Alignment of predicted tumor suppressing kinome reveals mutational hotspots in conserved regions

To define hot-spot inactivating missense mutations it was important to first establish a list of potential tumor suppressing kinases, identified by virtue of frequent truncation mutations. The top kinases identified could then be utilized to identify highly conserved residues that are mutated at a high frequency and will likely abolish catalytic activity, consistent with the kinase being a tumor suppressor. This screen was performed by locating the highly conserved APE motif or its equivalent sequence in 411 catalytically active (Supplementary Table S1) human kinases possessing the classical kinase domain motifs from Manning *et al* [16]. The APE motif, which is a critical component of the kinase domain functioning to stabilize the C-lobe and mediate substrate interactions, was used as a conservative cut-off point for identifying truncating mutations that would abolish kinase activity (Fig. 1). Although there are additional critical kinase regions C-terminal to this conservative cut off point, including the **a**F helix which functions to anchor both the catalytic and regulatory spines of the kinase domain, the APE motif was chosen because it is highly conserved and can be easily aligned across the kinase family. When adjusting the cut-off to include the entire protein, there are 11 unique kinases included in the top 30, therefore a majority of the top kinases would still be identified as tumor suppressing kinases (Fig. 2 and Supplementary Table S2). The frequency of truncating mutations found within the TCGA and CCLE datasets occurring N-terminal to this cut-off for each kinase were length corrected (Fig. 1) to produce a list of the top 30 kinases by truncating mutation density (Fig. 2 and Supplementary Table S3). This list is comprised of known tumor suppressors such as STK11 and MAP2K4 [20,21] along with kinases without a previously published role in tumor suppression. As a proof-of-concept, we verified one such MAP2K4 truncating mutation (E221*) found in a pancreatic cell line (CAPAN 1) to show that when wild type signaling is restored a significant decrease in anchorage-dependent and anchorage-independent colony forming potential is observed (Supplementary Fig. S1A-C). The kinase domains of these 30 tumor suppressing kinases were then sequence aligned to identify areas of high conservation. The TCGA and CCLE datasets were queried to capture all missense mutations occurring at each position of the aligned sequences and a combined score based on conservation and mutational frequency was produced for each position (Fig. 2 and Supplementary Table S4). The top 12 residues identified by this combined score (with a conservation score above 20) were located within a motif known to be critical for kinase activity (Table 1) [22, 23]. A total of five novel regions were identified within the top 20 hotspot residues (Table 1, highlighted with blue or grey). Two of these five residues (APE-6 and HRD-6) were further validated to determine their effect on kinase catalytic activity as discussed below.

**Figure 1.**
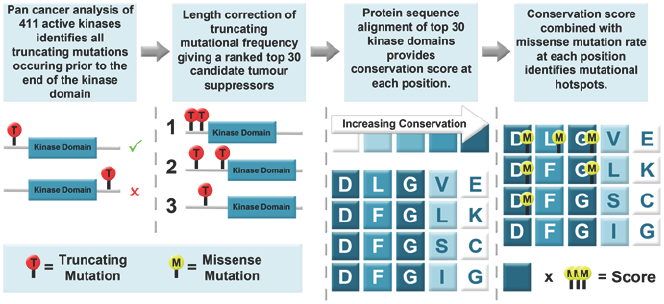
Schematic illustrating the screen of 411 kinases to produce the list of hotspot inactivating residues. A pan cancer analysis across TCGA and CCLE datasets was performed to identify truncating mutations occurring N-terminal to the APE motif. The truncation mutation frequency was length corrected to account for intra-kinase variability due to the position of the kinase domain within the overall protein. The kinase domains (GxGxxG to APE motif) of the top 30 kinases determined by length corrected truncation mutation frequency were sequenced aligned to identify conserved codons (between the 30 top kinases). This allowed re-querying of TCGA and CCLE datasets for mutational frequency at each residue. The conservation and mutational scores were combined to rank each residue of the kinase domain to generate a list of hotspot residues for kinase inactivating mutations.

**Figure 2.**
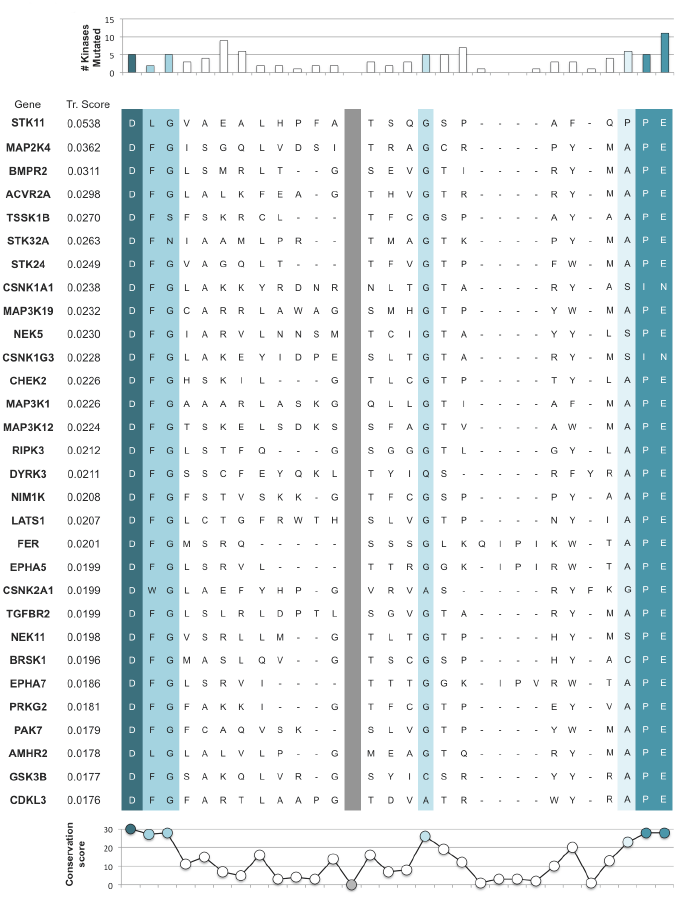
Output of the truncation mutation screen with the top 30 kinases illustrating a region of kinase sequence analyzed for conservation and number of kinases harboring missense mutations at each position. The top 30 kinases found from the truncation mutation screen, ranked by descending length corrected truncation mutation score (Tr. Score). A portion of the kinase sequence alignment (DFG to APE motif) is shown as an example to illustrate the tumor suppressing hotspot mutation screen. The vertical grey bar highlights a break in the alignment shown, as this region had very poor sequence alignment. The number of kinases mutated at each position is graphed along the top of this alignment whilst the conservation score for each residue is graphed along the bottom. The conservation score corresponds to the number of kinases with the most common amino acid at that position. Residues with high levels of conservation (>20) are colored in shades of blue. The full alignment performed from GxGxxG to APE is available in Supplementary Table S4.

**Table 1.**
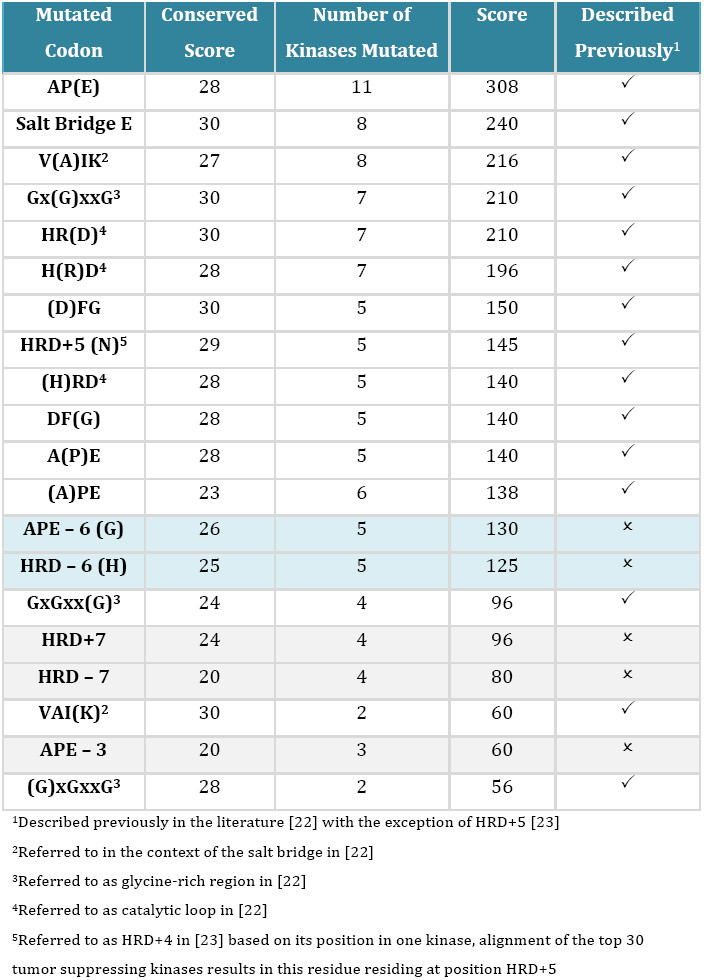
Tumor suppressing hotspot residues, identified through our motif screen, ranked by their total score (the product of the conservation and mutational scores). Note residues with conservation scores below 20 are excluded. Blue shading indicates validated new hotspot LOF residues, grey shading indicates new residues that are predicted to be LOF.

| Mutated Codon | Conserved Score | Number of Kinases Mutated | Score | Described Previously<sup>1</sup> |
| --- | --- | --- | --- | --- |
| AP(E) | 28 | 11 | 308 | ✓ |
| Salt Bridge E | 30 | 8 | 240 | ✓ |
| V(A)IK<sup>2</sup> | 27 | 8 | 216 | ✓ |
| Gx(G)xxG<sup>3</sup> | 30 | 7 | 210 | ✓ |
| HR(D)<sup>4</sup> | 30 | 7 | 210 | ✓ |
| H(R)D<sup>4</sup> | 28 | 7 | 196 | ✓ |
| (D)FG | 30 | 5 | 150 | ✓ |
| HRD+5 (N)<sup>5</sup> | 29 | 5 | 145 | ✓ |
| (H)RD<sup>4</sup> | 28 | 5 | 140 | ✓ |
| DF(G) | 28 | 5 | 140 | ✓ |
| A(P)E | 28 | 5 | 140 | ✓ |
| (A)PE | 23 | 6 | 138 | ✓ |
| APE - 6 (G) | 26 | 5 | 130 | × |
| HRD - 6 (H) | 25 | 5 | 125 | × |
| GxGxx(G)<sup>3</sup> | 24 | 4 | 96 | ✓ |
| HRD+7 | 24 | 4 | 96 | × |
| HRD - 7 | 20 | 4 | 80 | × |
| VAI(K)<sup>2</sup> | 30 | 2 | 60 | ✓ |
| APE - 3 | 20 | 3 | 60 | × |
| (G)xGxxG<sup>3</sup> | 28 | 2 | 56 | ✓ |

<sup>1</sup>Described previously in the literature [22] with the exception of HRD+5 [23]
<sup>2</sup>Referred to in the context of the salt bridge in [22]
<sup>3</sup>Referred to as glycine-rich region in [22]
<sup>4</sup>Referred to as catalytic loop in [22]
<sup>5</sup>Referred to as HRD+4 in [23] based on its position in one kinase, alignment of the top 30 tumor suppressing kinases results in this residue residing at position HRD+5

### Mutations of a hinge residue between the activation and P+l loops abolish catalytic function

The APE-6 residue is a glycine residue found to be highly conserved within the 411 kinases used in this study (81% conservation across 411 kinases). Structural modelling demonstrates that this residue lies at a hinge point between the activation loop and P+1 loop which could allow the typical fluctuations of the activation loop between its active and inactive conformations (Fig. 3A). The small size of the glycine amino acid that occupies this position in a large number of kinases allows for the flexibility of this hinge region. Mutations that limit flexibility of the hinge region may impair catalytic activity. Alternatively, this glycine helps form the P+1 recognition pocket and mutations at this residue will impact this pocket to alter substrate recognition and binding, which could also lead to LOF. Molecular dynamics (MD) simulations of one such mutation at this conserved glycine in MAP2K4 (G265D) showed a decreased movement of the activation loop compared to that of the wild type kinase (Fig. 3B). Transient overexpression of MAP2K4 mutations, G265D and G265C, both seen in cancer samples, demonstrates reduced phosphorylation of the canonical JNK pathway equivalent to a kinase-dead construct (Fig. 3C). When wild type MAP2K4 was stably re-expressed in CAL51 cells, harboring the G265D mutation (Supplementary Fig. S2D), a reduction in colony forming potential was observed in both two-dimensional and three-dimensional assays (Fig. 3D and E), compared to the parental cell line, where no significant change was observed. These data indicate that the G265D MAP2K4 mutation is a significant LOF driver mutation in the CAL51 cell line. Corresponding glycine mutations observed in other cancer samples in MAP3K13 (G315D) and PRKCQ (G541V) also showed a reduction in kinase activity (Fig. 3F and G).

**Figure 3.**
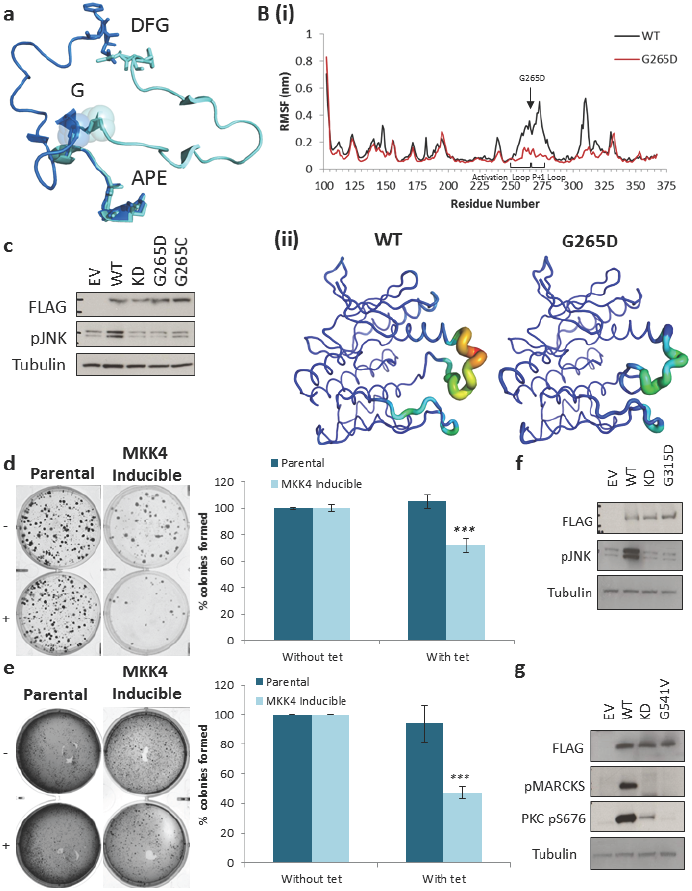
Mutations identified in a hinge glycine at position APE-6 are inactivating. A) Position of the conserved APE-6 glycine at a hinge point of the activation loop (shown in INSR kinase domain). Active kinase conformation is shown in light blue (PDB ID: 1IR3), inactive kinase conformation is shown in dark blue (PDB ID: 1IRK). DFG and APE motifs are shown as sticks, glycine is shown in sticks and spheres. B) MD simulations show a large decrease in the level of movement observed for the activation loop in MAP2K4 (PDB ID: 3ALN). (i) Root-mean-squared fluctuations (RMSF) of each residue is shown graphically and (ii) structurally, with width and color of the ribbon showing corresponding level of movement. C) Western blot showing biochemically that mutations in the conserved glycine of MAP2K4 are LOF towards to the JNK pathway. A significant decrease in 2D (D) and 3D (matrigel 3D embedded growth assay); E) colony forming potential is observed in CAL51 cell line harboring MAP2K4 G265D following tetracycline-inducible expression of wild type MAP2K4 (*** = *p <* 0.001). Mutations of the conserved APE-7 glycine within MAP3K13 (F) and PKC6(G) are also LOF. Error bars depict the standard error of the mean; statistical significance was calculated with a two-tailed Students t-test.

### Mutations at HRD-6 abrogates kinase activity

Investigation into the HRD-6 position was also performed. Structural modelling of this residue highlights its close proximity to the R-spine anchoring residue within the aF-helix (Fig. 4A). The R-spine is formed as the kinase becomes active and is critical for catalytic activity. MD simulation of a mutation at the HRD-6 position within DAPK3 (H131R) shows an increased level of movement around three out of the four R-spine residues, with RS1 at residue position 79 showing a large increase in movement. This increased movement could suggest a destabilization of the R-spine, which would likely result in reduced catalytic activity. Assessment of cancer associated mutations at this position in DAPK3 (H131R) and BRAF (H568D) demonstrate a loss of kinase catalytic activity similar to that observed for kinase dead mutants (Fig. 4B and C). These results highlight that combining conservation at a given residue with mutational frequency will be a successful approach to identify low frequency functional mutations across cancer genomic databases.

**Figure 4.**
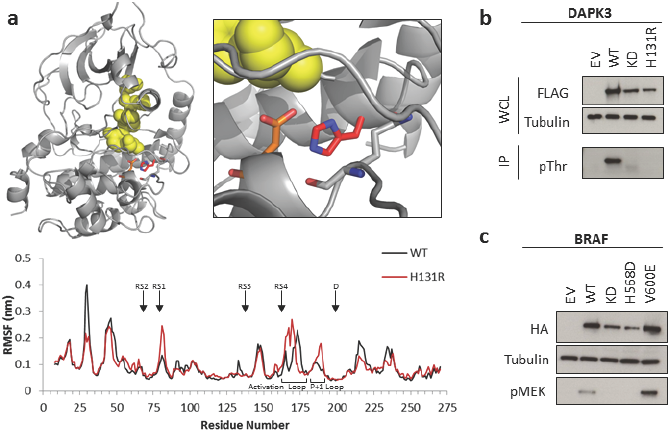
Mutations identified in HRD-6 are inactivating. A) HRD-6 residue (red sticks) lies in close proximity to the R-spine anchoring residue within **α**F helix (orange sticks). R-spine shown as yellow spheres (PKA-PDB ID: 1ATP). Root-mean-squared fluctuations (RMSF) from MD simulations highlight increased movement around R spine residues RSI, Rs3 and RS4 in DAPK3 H131R (PDB ID: 3BHY), which results in an altered movement of the activation and P+l loops. B) In vitro kinase assay shows decreased kinase activity within DAPK3 H131R as determined by decreased autophosphorylation. C) Decreased kinase activity observed for BRAF H568D mutation, as observed by downstream phosphorylation of MEK.

### Pan cancer analysis of mutations found in critical motifs highlights a prevalence of LOF MAP2K7 mutations in gastric cancer

Having identified and validated novel hotspot residues for kinase inactivation, the top 15 mutational hotspots (13 known critical residue and 2 novel residues validated above) were used in a functional screen to identify novel tumor suppressing kinases. A pan cancer analysis was performed by querying the TCGA and CCLE datasets for mutations located in these 15 residues in 411 kinases. The kinases were then ranked by the mutational frequency of these 15 regions combined (Fig. 5A). BRAF was identified as the top hit with 39 mutations throughout both datasets, followed by STK11 (14 mutations), MY03A (13 mutations), EPHB1 (12 mutations), and MAP2K7 (11 mutations). EPHB1 and STK11, as well as other top hits such as CHEK2, have previously been demonstrated to play a tumor suppressive role in different cancer sub-types [21, 24–26]. MAP2K7 was selected for further investigation as 6 out of the 11 detected mutations occurred in a single cancer subtype, gastric adenocarcinoma (Fig. 5B). In addition, MAP2K7 mutations and deletions occur in 7% of cases in the TCGA gastric adenocarcinoma series, with 40% of the mutated cases possessing more than one mutation, suggesting both alleles are affected. Transient overexpression of the 4 gastric MAP2K7 mutations that occurred in conserved motifs validated that three of these mutations are LOF with regards to JNK pathway activation (Fig. 5C). Furthermore, in a gastric cancer cell line that harbors an inactivating MAP2K7 mutation (IM95, D290fs, Supplementary Fig. S2A), reconstitution of wild type signaling (Supplementary Fig. S2C) results in a significant decrease in anchorage-dependent and anchorage-independent colony forming potential (Fig. 5D and E).

**Figure 5.**
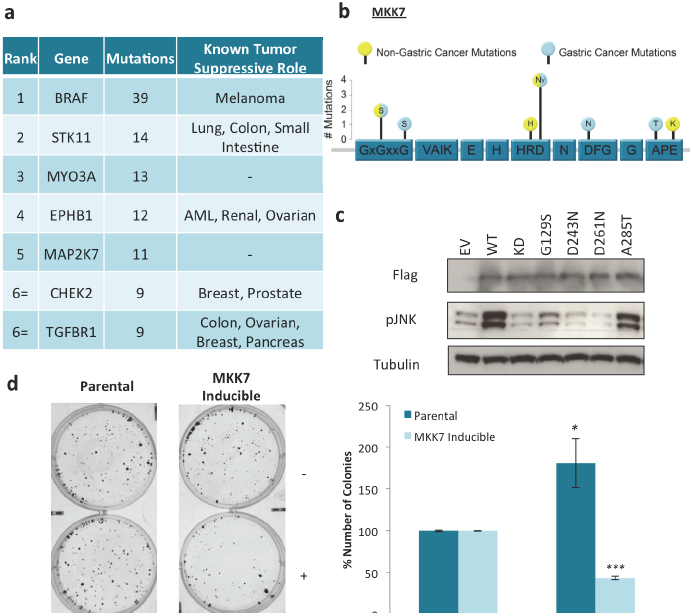
Critical codon screen identifies MAP2K7 as a target in gastric cancer. A) Table ranking kinases based on the number of mutations observed within top 15 critical codons (Table 1). B) Schematic highlighting mutations identified within key motifs of MAP2K7. Mutations are indicated by yellow spheres, those colored blue are found in gastric cancer, those colored half blue have half of the mutations observed in gastric cancer. C) Western blot showing MAP2K7 gastric cancer mutations are LOF towards downstream substrate JNK. D) 2D colony formation assay with IM95 gastric cell line harboring LOF MAP2K7. Parental cell line shows increased numbers of colonies formed following addition of tetracycline. IM95 cell line with tetracycline-inducible expression of MAP2K7 (IM95+MAP2K7) shows significantly decreased numbers of colonies formed following the addition of tetracycline (*** = *p <* 0.001, * = *p <* 0.05). E) 3D colony formation assay with the IM95 cell lines. IM95+MAP2K7 cell line shows significantly decreased colony forming potential when MAP2K7 signaling is restored following tetracycline treatment. No difference is observed when the parental cells are treated with tetracycline (*** = *p <* 0.001). Error bars depict the standard error of the mean; statistical significance was calculated with a two-tailed Students t-test.

When other genes in the JNK pathway are considered, approximately 22% of gastric cancers harbor alterations in MAP2K7, MAP2K4, MAPK8, MAPK9, MAPK10, JUN, or ATF2 with a high degree of mutual exclusivity suggesting a significant role for loss-of-function on the JNK pathway in gastric carcinogenesis (Supplementary Fig. S2D). Re-expression of wild type JNK1 in a cell line harboring a JNK1 LOF mutation (G177*; Supplementary Fig. S2B-C) also shows significant decrease in anchorage-dependent growth (Supplementary Fig. S1E). Together these data indicate that loss of signaling through the JNK pathway is important for gastric cancer tumorigenesis.

## DISCUSSION

Cancer genomics has provided large cohorts of data which has proved extremely valuable in identifying common genetic drivers. However, the challenge now lies in dissecting this data to identify those driver mutations that are at a lower frequency, and thus masked by mutational noise. We present an approach to screen genomic data utilizing functional knowledge of the kinase domain structure and sequence. Our first step was to utilize a strategy to identify somatic point mutations that will abolish the catalytic activity of a kinase, and through a pan cancer screen, identify potential tumor suppressing kinases enriched in LOF mutations. Following this filtering step, we identified a number of kinases, such as MAP2K7, that we would predict to harbor frequent LOF mutations. Our final step was to validate our approach by demonstrating biochemically and through functional assays that indeed MAP2K7 is a tumor suppressing kinase in gastric cancer. To put this challenge in perspective, at the time of conducting our screen, there were 43,212 missense mutations in the 411 canonical kinases included in our screen. From our conserved motif screen, that included 15 highly conserved and mutated codons, we identified 921 mutations in these 411 kinases, allowing us to pinpoint novel kinases enriched in LOF mutations. Our strategy has not only highlighted many additional kinases to be explored as tumor suppressors, but also laid the ground work for a broadly applicable approach that can be utilized to identify novel tumor suppressors in other enzyme families, such ubiquitin ligases.

The initial truncation screen identified a list of candidate tumor suppressing kinases and to ensure that the truncation mutations used within this screen would result in kinase inactivation a conservative cutoff point for these truncation mutations was determined. We only included truncating mutations occurring N-terminal to the APE motif, as we could confidently state these would abolish kinase activity. The length-corrected truncation mutation frequency was used to rank all the kinases as potential tumor suppressors. The appearance of two well-known tumor suppressing kinases (STK11 and MAP2K4) occurring as the top two hits for our screen validated this approach [20,21].

To identify novel hotspot tumor suppressive residues we aligned the sequences of the top 30 tumor suppressing kinases and each residue was assigned a mutational‐ and conservational-score. Residues were then ranked by their combined score to give a list of tumor suppressive hotspot residues. The top 12 residues identified in this way occurred in well-known motifs critical for kinase function, validating this approach as a method for identifying highly functional mutations. However, we accept that by virtue of their conservation and prior recognition there is potential bias in the sequence alignment, which would lead to a higher score for these well-known motifs. We investigated the two novel residues that occurred in the top 15 hotspot residues identified by this screen. The first of these was a conserved glycine, occurring between the activation and p+1 loops, which is frequently mutated in different kinases in cancer samples. We demonstrated that mutation at this flexible hinge residue reduced the movement of the activation loop resulting in kinase inactivation, highlighting this residue as critical for kinase function. The other novel residue to be ranked within the top 15 hotspot residues was a highly conserved histidine residue at the HRD-6 position. This residue is located in close proximity to the R-spine anchor residue within the **a**F helix, known to be critical for stability of this R-spine [27]. MD simulations highlighted increased instability in the R-spine residues, which correlated to the loss of kinase activity in a number of different kinases

Having identified 15 residues within the kinase domain that were either known to be critical to kinase function, or validated to result in a LOF phenotype when mutated, a screen to identify kinases that harbor lower frequency genetic drivers was performed. Due to the small target area being screened in each sample a large dataset was required; therefore a pan-cancer analysis was performed utilizing the TCGA and CCLE datasets. This analysis identified any mutations occurring in these 15 residues in 411 kinases across all cancer sub-types. Like the truncation mutation screen this approach identified known tumor suppressors, many of which were common to both screens (such as STK11, CHEK2 and TGFBR1). Given that the critical codon screen was derived from the truncation screen data some overlap could be expected. However, there were also marked differences between the results of the two screens with the oncogene BRAF featuring as the top kinase in the critical motif screen whilst only ranking 88^th^ out of 411 kinases in the truncation screen. Kinase-dead BRAF can paradoxically act as a scaffold to activate CRAF resulting in activation of the MEK/ERK pathway [24]. It is likely that the majority of truncation mutations interfere with this oncogenic mechanism resulting in fewer truncation mutations being observed in the BRAF oncogene. With this consideration in mind, comparing the results of our two screens may help to identify mechanisms by which kinase dead mutations paradoxically activate signaling pathways by a similar mechanism, rather than resulting in loss of function of the canonical signaling pathway. Therefore, the discrepancies that exist for the two tumor suppressing lists can shed light on exciting biology in regards to inactivating mutations that activate a pathway.

Finally, the screens we developed here identified MAP2K7, an upstream activator of the JNK pathway, as one of the top hits when identifying mutations in codons critical for kinase function. Interestingly, the majority of these mutations (6/11) were observed in gastric cancer samples highlighting inactivating MAP2K7 mutations as gastric cancer drivers. Furthermore, when genomic data from the cBioPortal was interrogated, other JNK pathway components, including JNK, JUN and ATF, were observed to be mutated or deleted in gastric cancer. The JNK pathway dictates key processes regulating cancer development but its role in tumorigenesis remains controversial with research showing it acting as both a pro-oncogene or tumor suppressor [reviewed in 26]. This dual nature of the JNK signaling pathway has been shown to be highly dependent on the cellular context and extracellular environment [19]. For example, decreasing levels of JNK in a background of oncogenic Ras or reduced p53 enhances lung carcinogenesis and mammary tumor development respectively [18]. Conversely, JNK inactivation has been shown to result in reduced hepatocellular carcinoma development and a decrease in tumors in a carcinogen induced gastric cancer model [18, 29]. Disparate roles for JNK signaling within the same cancer types are also observed. In the context of colon cancer, JNK-mediated cJUN phosphorylation was shown to be crucial for intestinal cancer development in an APC dependent colorectal cancer mouse model (APCMin/+ mice) with inactivation of cJUN resulting in fewer, smaller tumors [30]. Additionally, it has been shown that increased JNK activity accelerated colorectal tumorigenesis through direct cross-talk with Wnt signaling pathway components, resulting in an oncogenic positivity feedback loop [31]. Whilst, the loss of JNK-mediated upregulation of p21 resulted in the spontaneous development of intestinal tumors in JNKl-deficientmice [32] and increased proliferation in a human colorectal cancer cell line [33]. This contrasting role is also observed for MAP2K7 with high levels of MAP2K7 showing enhanced proliferation and metastasis in lung, colon, hepatocellular, prostate and breast cancers [34–38]. Conversely, MAP2K7 has also been shown to slow the progression of lung and mammary tumors driven by other oncogenic drivers [39–40]. Interestingly, our tumor suppressive screens also identified LOF mutations in MAP2K4 and MAP3K13, upstream JNK pathway activators. Our experimental data highlights that inactivation of the JNK pathway is important for promoting tumorigenic phenotypes in gastric cancer, which offers insight into a molecular mechanism that could affect up to 22% of gastric cancer patients. Furthermore, our genetic data highlighting MAP2K7, MAP2K4 and MAP3K13 mutations in different cancer subtypes could suggest that loss of signaling in the JNK pathway promotes tumorigenesis in a number of human tumor types

Gastric cancers carry a high number of mutations per sample although few specific driver mutations are known [41]. By focusing on mutations with a high probability of disrupting kinase function our screen helps remove mutational noise caused by passenger mutations. Using the MAP2K7 observation as a prompt to query other known pathway members highlights a prominent role for JNK pathway inactivation in almost a quarter of all gastric cancers. It is proposed that sequencing more cancer samples will eventually cause lower frequency drivers to become more apparent [42]. Whilst this argument may be true it is clear from our experience with analysis of the TCGA gastric cancer dataset, where 287 cancers have been fully sequenced and over 1000 genes have a mutational frequency over 5%, that this process is greatly facilitated when precise functionally derived algorithms are integrated into the pipeline [9].

In conclusion, we have developed and validated two approaches that filter freely available NGS data with functional consideration to identify novel genetic drivers that would otherwise be lost within the mutational noise. Our first approach utilized truncating mutation frequency to identify tumor suppressive kinases and, from these, identified novel residues critical for kinase function. Utilizing these critical codons we identified kinases harboring high numbers of LOF mutations, revealing more tumor suppressing kinases with additional information on tumor type enrichment. Taken together these screens not only provide novel tumor suppressors to investigate, but also highlight novel residues of critical importance in kinase activation. This approach, utilizing the human kinome, provides a wealth of information on inactivated kinases in cancer, and could be expanded to identify both GOF and LOF mutations within other proteins containing conserved protein domains.

## METHODS

### Truncation mutation screen

R scripts were prepared to identify all kinase truncating mutations within the TCGA (Script1) and CCLE (Script2) databases occurring before the end of the kinase domain. In brief, the APE motif was identified in the Genbank sequences of 411 catalytically active kinases with conventional VAIK, HRD and DFG motifs identified in Manning *et al.* (Supplementary Table S1) [16]. The location of the E of the APE motif was defined as the C-terminal limit of a functional kinase domain. Mutational data from TCGA and CCLE were cross referenced with each kinase APE location to record all truncating mutations occurring N-terminal to this location. Mutational frequencies were length corrected using the mean transcript length (between shortest and longest transcripts) of each kinase to the E of the APE motif to account for inter-kinase differences. Supplementary Table s3 also includes alternate top 30 kinases based on alteration of the cut-off at APE to include less conservative limit including using the whole protein. A top 30 tumor suppressing kinase list was constructed by ranking kinases by descending length corrected score (Figures 2, S3 and Supplementary Table S3) and the kinase domain sequences of these 30 kinases was identified (Script3). Where a kinase was annotated as possessing more than one kinase domain the domain sequence with the motif configuration most closely resembling a classical kinase domain was selected for analysis.

### Sequence alignment and residue conservation and mutation scoring

The kinase domain sequence from the first glycine of the GxGxxG loop to APE motif were sequence aligned for each of the top 30 kinases using the Strap Alignment Tool. The conservation score was determined as the number of kinases out of the top 30 that harbored the most common amino acid at that position. If a location had more than 5 kinases without a corresponding amino acid (in other words this region is missing from 5 or more kinases) the conservation score was calculated as zero. All loci with a conservation score below 20 were removed from the final analysis to leave highly conserved locations. The aligned sequence locations were cross-referenced with the mutational data from TCGA and CCLE to identify the mutations at each position and the mutational score was calculated as the number of kinases with a mutation at that position. Multiplying the mutational score by the conservation score produced a combined score to rank each residue in the kinase domain sequence. Supplementary Table S4 displays the scores and alignment for all residues with at least one mutation and Supplementary Figure S4 displays the frequency distribution of combined scores for each residue with a least one mutation.

### Kinase motif based screen

R scripts were written to identify any mutations from the TCGA (Script5) and CCLE (Script6) databases occurring within the hotspot residues identified above. In brief, the top 15 hotspot residues (defined by combined score above) were located within the Genbank sequences for each kinase. Mutational data from the TCGA studies was obtained through the CGDS-R package (Supplementary Table S5) and CCLE were cross-referenced with each kinase to identify any point mutations occurring within critical kinase motifs. A ranked list was constructed from the number of mutations observed per kinase (Script 7).

### Structural modelling and molecular dynamics simulations

Homology models of wild type and mutant MAP2K4 were created using Modeller 9.16 from PDB ID: 3ALN. Molecular dynamics simulations were performed using GROMACS version 5.0 with the GROMOS96 53a6 force field parameter set. All titratable amino acids were assigned their canonical state at physiological pH, short-range interactions were cut off at 1.4 nm and long-range electrostatics were calculated using the particle mesh Ewald summation [43]. Dispersion correction was applied to energy and pressure terms accounting for truncation of van der Waals forces and periodic boundary conditions were applied in all directions. Protein constructs were placed in a cubic box of 100 nM NaCl in simple point charge water with at least 1 nm distance between the protein construct and box edge in all directions. Neutralizing counter ions were added and steepest decent energy minimization was performed, followed by a two-step NVT/NPT equilibration. Both equilibration steps maintained a constant number of particles and temperature, NVT equilibration was performed for 100 ps maintaining a constant volume, followed by 10 ns of NPT equilibration maintaining a constant pressure. Temperature was maintained at 37°C by coupling protein and non-protein atoms to separate temperature coupling baths [44], pressure was maintained at 1.0 bar (weak coupling). All position restraints were then removed and simulations were performed for 400 ns using the Nose-Hoover thermostat [45] and the Parrinello-Rahman barostat [46]. Root-meansquared fluctuation (RMSF) analysis compared the standard deviation of the atomic position of each a-carbon in the trajectory, fitting to the starting structure as a reference frame. Root-mean-squared deviation (RMSD) analysis compared the structure of specified groups of residues at each time point of a trajectory with the reference starting structure. Images were created using PyMol version 1.5.0.5.

### Plasmids and transfections

BRAF and MAP3K13 cDNA were prepared from RNA extracted from HEK293T cells. PRKCQ was bought in the pENTR vector (Ultimate Human ORF Library ‐Life Technologies), MAP2K4 and MAP2K7 were bought in pCMV6-Entry vectors (Origene), and MLK4 was purchased in pReceiver-M12 FLAG vector (GeneCopoeia). Primers containing attB flanking sites were used to amplify up the constructs before they were inserted into pDONR-221 vector using the BP clonase reaction. The ABL1 constructs were purchased in the pDONR-223 vector. From here, the Gateway system was used for cloning into a pDEST-FLAG vector created by Dr. Eleanor Trotter from the pReceiver-M12 plasmid (GeneCopoeia). 3X-FLAG DAPK3 vector was provided by Dr. Timothy Haystead (Department of Pharmacology and Cancer Biology, Duke University Medical Center, Durham, North Carolina 27710).

Mutants were created using the Quikchange Site-directed Mutagenesis II Kit (Agilent Technologies) using the manufacturers protocol. Kinase dead mutants were BRAF (K483M), DAPK3 (K42M), MAP2K4 (K131M), MAP2K7 (K148M), MAP3K13 (K195M), MLK4 (K151M) and PRKCQ (K409M). All sequences were confirmed using Sanger sequencing. HEK293T cells or CAL51 cells (for MAP2K4 transfections) were seeded into 12-well plates (standard transfections) or 6-well plates (immunoprecipitations) and transiently transfected the following day using either Attractene (QIAGEN) for the HEK293T cells or Lipofectamine2000 (Thermo Fisher Scientific) for CAL51 cells according to the manufacturers protocol.

### Protein lysate preparation and immunoblots

Cells were lysed on ice after 24 hours using Triton X-100 Cell Lysis Buffer (Cell Signaling) supplemented with a protease inhibitor tablet (Roche). Lysates were either resolved on SDS-PAGE gels followed by western blotting or used in an in vitro kinase assay (details below). Primary antibodies used were as follows: Flag M2 and a-tubulin (Sigma); MAP2K4, MAP2K7, pJNK (T183/Y185), pMARCKS (S152/S156), pPKC (S676), pThr, and p-cjun (S73) (Cell Signaling Technology).

Mouse or rabbit HRP-conjugated secondary antibodies were used (Cell Signaling Technologies). All western blots are representative of at least three independent experiments.

### In vitro kinase assays

Cell lysates from DAPK3 transfections were incubated with anti-Flag M2 affinity gel (Sigma) for at least 2 hours. Beads were washed with lysis buffer and kinase buffer (Cell Signaling Technology) and a kinase assay was performed in the presence of 200 μM ATP at 30˚C for 30 min to assess autophosphorylation. Following addition of 4x reduced SDS sample buffer, proteins were resolved by SDS-PAGE and analyzed by western blotting.

### Generation of MAP2K4, MAP2K7 and JNK1 tetracycline-inducible cell lines

Parental CAL51 (MAP2K4), CAPAN1 (MAP2K4), IM95 (MAP2K7) and NUGC3 (JNK1) were used to generate cells with tetracycline-inducible expression of wild type plasmids (cloned into pLenti/TO/V5-DEST vector) and pLenti3.3/TR (for tetracycline repressor expression) were transfected into 293FT cells using Lipofectamine2000 to generate a lentiviral stock. Cells were transduced with lentiviral stocks and cell lines generated by antibiotic selection (Blasticidin (Invitrogen) and Geneticin (Gibco)). Tetracycline (Invitrogen) was used to induce expression of wild type MAP2K4 (CAL51, CAPAN1), wild type MAP2K7 (IM95) or wild type JNK1 (NUGC3).

### Anchorage-dependent colony formation assay

Cells were seeded at approximately 100 cells/well in a 6-well plate format. The following day, tetracycline was added and cells were left to grow for 3 weeks with media changed every 2-3 days. Colonies formed and were fixed with ice-cold methanol, stained with 0.5% crystal violet (Sigma) solution made up in 25% methanol. Wells were thoroughly washed and air dried. For quantification, 2 ml of 10% acetic acid was added to each well, incubated for 20 mins with shaking and absorbance values read at 595 nm.

### Anchorage-independent colony formation assay

Anchorage-independent colony formation assays were performed with CAPAN 1, NUGC3 and IM95 cell lines. Plates were initially coated with 0.6% softagar with or without tetracycline containing no cells. Cells were then seeded at approximately 10,000 cells/well in a 6-well plate format in 0.35% soft agar with or without tetracycline. Media was added (with or without tetracycline) and changed every 2-3 days. After 3 weeks cells were stained using 0.05% crystal violet (Sigma) solution made up in 25% methanol.

### Matrigel 3D embedded growth assay

Matrigel 3D embedded growth assays were performed with the CAL51 cell line, as these cells did not grow successfully in the anchorage independent growth assay. 6 well plates were pre-coated with a thin layer of EHS (Corning) before cells were seeded at approximately 50,000 cells/well in EHS. 2 ml of media with or without tetracycline was added. Media was changed every 2-3 days and after 10 days cells were stained using 0.05% crystal violet (Sigma) solution made up in 25% methanol.

### Statistical Analyses

All statistical significance values were calculated using a two-tailed Students ttest.

## SUPPLEMENTARY MATERIALS

**Supplementary Figure S1: Truncation mutation screen identifies tumor suppressing kinases.** Stable tetracycline-inducible re-expression of WT MAP2K4 in the pancreatic cell line CAPAN1, which harbors a truncating MAP2K4 mutation, results in significant loss of anchorage-dependent (A) and anchorage-independent (B) colony forming potential (*** = *p* < 0.001, * = *p* < 0.05). Stable tetracycline-inducible cell lines CAPAN1 (C) and CAL51 (D) over-express WT MAP2K4 following the addition of tetracycline.

**Supplementary Figure s2. MAP2K7 pathway components offer a significant target for gastric cancer therapy.** A) MAP2K7 D290fs mutation observed in the IM95 gastric cancer cell line is inactivating towards downstream substrate JNK. B) JNK1 G177* mutation observed in the NUGC3 gastric cancer cell line is inactivating towards downstream substrate cjun. C) Stable tetracycline-inducible cell lines over expressing WT MAP2K7 (IM95) or JNK1 (NUGC3) following the addition of tetracycline. D) MAP2K7 pathway components are altered in 22% of patient samples sequenced within the TCGA and CCLE databases, data taken from the cBioPortal. (E) Re-expression of WT JNK1 in NUGC3 cell line harboring JNK1 G177* mutation shows a significant reduction in anchorage-dependent colony forming potential (*** = *p <* 0.001). targ et for gastric cancer therapy.

**Supplementary Figure S3:** Frequency histogram of truncation mutation scores (calculated as number of truncation mutations divided by mean length of gene upstream of APE motif). The Top 30 kinases are shaded blue to demonstrate that they are located in the tail of the distribution.

**Supplementary Figure S4:** Frequency histogram of combined scores for each codon in the kinase domain of Top 30 kinases (calculated as mutation score multiplied by conservation score). The top 15 codons based on score (in addition to two codons with high scores based on mutation score but with a lower conservation score than the cut-off of 20 to make the final list) are shaded blue to demonstrate that they represent the tail of the distribution. Note that this distribution does not include residues with a combined score of 0 (as a result of a mutational score of 0).

**Supplementary Table SI:** List of 411 active kinases used in the screens within this paper.

**Supplementary Table s2:** Table displaying the effect on the top 30 candidate tumour suppressors by changing the cut-off point used to determine truncating mutations that abolish kinase catalytic activity. The APE columns display the original screen used in the manuscript with the cut-off being the E of the APE motif. APE+50 and APE+100 display the top 30 when the cut-off is extended past the E of the APE motif by 50 or 100 residues respectively. The total column displays the top 30 truncation mutations when the whole length of the gene is used. The grey shaded kinases are those that do not appear in the top 30 when the respective cut-offs are used and the kinases in bold are those that appear in the new list.

**Supplementary Table S3:** Truncation mutation scores for all 411 kinases. As numerous gene transcripts exist, gene length was calculated by taking an average of the longest and shortest transcript lengths from the start of the gene to the codon encoding the E of the APE motif.

**Supplementary Table S4:** Output from the computational screen aligning the top 30 tumour suppressing kinases, identified in an initial screen, and ranking each amino acid within the kinase domain based on their conservation and mutational scores (for all residues with at least 1 mutation).

**Supplementary Table S5**: TCGA studies from which mutational data was acquired on 7th January 2016.

## Other Supplementary Information

Script1_TruncatingMutations_TCGA.R

Script2_TruncatingMutations_CCLE.R

Script3_Truncating Mutations_Merge CCLE and TCGA data.R

Script4_Identification of Motif Residues.R

Script5_Genbank Merge with TCGA.R

Script6_Genbank Merge with CCLE.R

Script7_Merge TCGA and CCLE.R

Script Source files

## ACKNOWLEDGEMENTS

We would like to thank Professor Alexandra Newton and Dr. Timothy Haystead for their kind donation of some of the DNA constructs used in this study.

## Author contributions

AH, NLS and JB designed research; AH, NLS and AJF wrote the computational script; NLS performed the computational simulations; NLS, CL, ET, GK, PBK and MH performed the experiments; AH, NLS, ET, CW, CM and JB analyzed the data; AH, NLS and JB wrote the manuscript.

## Data and materials availability

All DNA plasmids mentioned in this paper will be made available to the research community either through AddGene or direct requests to the laboratory. Scripts and data files utilized in the computational analysis can be accessed through the GitHub repository found at https://github.com/adamfletcherUK/Truncation_and_motif_based_analysis.git.

